# Drosophila ring chromosomes interact with sisters and homologs to produce anaphase bridges in mitosis

**DOI:** 10.1101/2024.08.08.607186

**Authors:** Ho-Chen Lin, Mary M. Golic, Hunter J. Hill, Katherine F. Lemons, Truc T. Vuong, Madison Smith, Forrest Golic, Kent G. Golic

## Abstract

Ring chromosomes are known in many eukaryotic organisms, including humans. They are typically associated with a variety of maladies, including abnormal development and lethality. Underlying these phenotypes are anaphase chromatin bridges that can lead to chromosome loss, nondisjunction and breakage. By cytological examination of ring chromosomes in *Drosophila melanogaster* we identified five causes for anaphase bridges produced by ring chromosomes. Catenation of sister chromatids is the most common cause and these bridges frequently resolve during anaphase, presumably by the action of topoisomerase II. Sister chromatid exchange and chromosome breakage followed by sister chromatid union also produce anaphase bridges. Mitotic recombination with the homolog was rare, but was another route to generation of anaphase bridges. Most surprising, was the discovery of homolog capture, where the ring chromosome was connected to its linear homolog in anaphase. We hypothesize that this is a remnant of mitotic pairing and that the linear chromosome is connected to the ring by multiple wraps produced through the action of topoisomerase II during establishment of homolog pairing. In support, we showed that in a ring/ring homozygote the two rings are frequently catenated in mitotic metaphase, a configuration that requires breaking and rejoining of at least one chromosome.

## Introduction

Eukaryotic genomes are typically organized into sets of linear chromosomes, each representing a single DNA molecule bounded by telomeres at each end. However, circular, or ring, chromosomes have been found as abnormal variants in many eukaryotic species. Eukaryotic ring chromosomes were first identified by L.V. Morgan in *Drosophila melanogaster* and soon thereafter by McClintock in corn (Morgan 1926; McClintock 1932). Ring chromosomes have since been found, or constructed, in *Saccharomyces cerevisiae*, *Schizosaccharomyces pombe, Caenorhabditis elegans*, humans, cattle, dogs, cats, rats, mice, Chinese hamsters, a variety of plant species, and likely other examples that we have overlooked (Ramiah et al. 1935; Tsunewaki 1959; Lindsten and Tillinger 1962; Turner et al. 1962; Wang et al. 1962; Kikuchi et al. 1979; Strathern et al. 1979; Morgan et al. 1986; Bartnitzke et al. 1992; Fan et al. 1992; Voet et al. 2003; M. et al. 2011; Murata et al. 2013; Murata 2014; Szczerbal et al. 2017; Koo et al. 2018; Wang et al. 2018; Rappaport et al. 2021). They may arise spontaneously by homologous or nonhomologous recombination between sequences on opposite sides of a centromere, by telomere dysfunction and fusion of opposite ends of a chromosome, by centromere misdivision combined with an additional break, or deliberately constructed through a variety of genetic manipulations. Ring chromosomes can be found in human cancers and are typically predictive of poor patient outcomes (Levan 1956; Gebhart 2008). Constitutional ring chromosomes in humans are associated with developmental and intellectual disabilities (Kosztolányi 2012; Yip 2015).

Ring chromosomes also exhibit aberrant behaviors, such as mitotic instability and dominant lethality. Cytological analyses in a variety of organisms have shown anaphase chromatin bridges and the occurrence of rings with varied sizes in a single individual (Morgan 1926; McClintock 1932; Morgan 1933; McClintock 1938)(Braver and Blount 1950; Hinton 1955; Levan 1956; Hinton 1959; Pasztor 1971; Stone 1982). There is some uncertainty about the precise origin and nature of anaphase bridges, and the degree to which these may vary between organisms and between different chromosomes. Anaphase bridge formation has been attributed to problems associated with the segregation of catenated rings, to dicentric chromosomes generated by sister chromatid exchange (SCE), or to chromosome breakage and joining of sister chromatids to make a foldback union (SCU; Fig. 1A-C) (McClintock 1938; Braver and Blount 1950; Hinton 1955; Levan 1956; Hinton 1957; Hinton 1959; Brosseau 1966). Variation in ring chromosome number and size can be attributed to nondisjunction or breakage of chromosomes involved in bridges. Because these reports on ring chromosomes lack photographic documentation of mitotic behaviors it seemed worthwhile to carry out a detailed examination of ring behavior, and to include not only ring-*X* chromosomes, but ring-autosomes as well.

**Figure 1:**
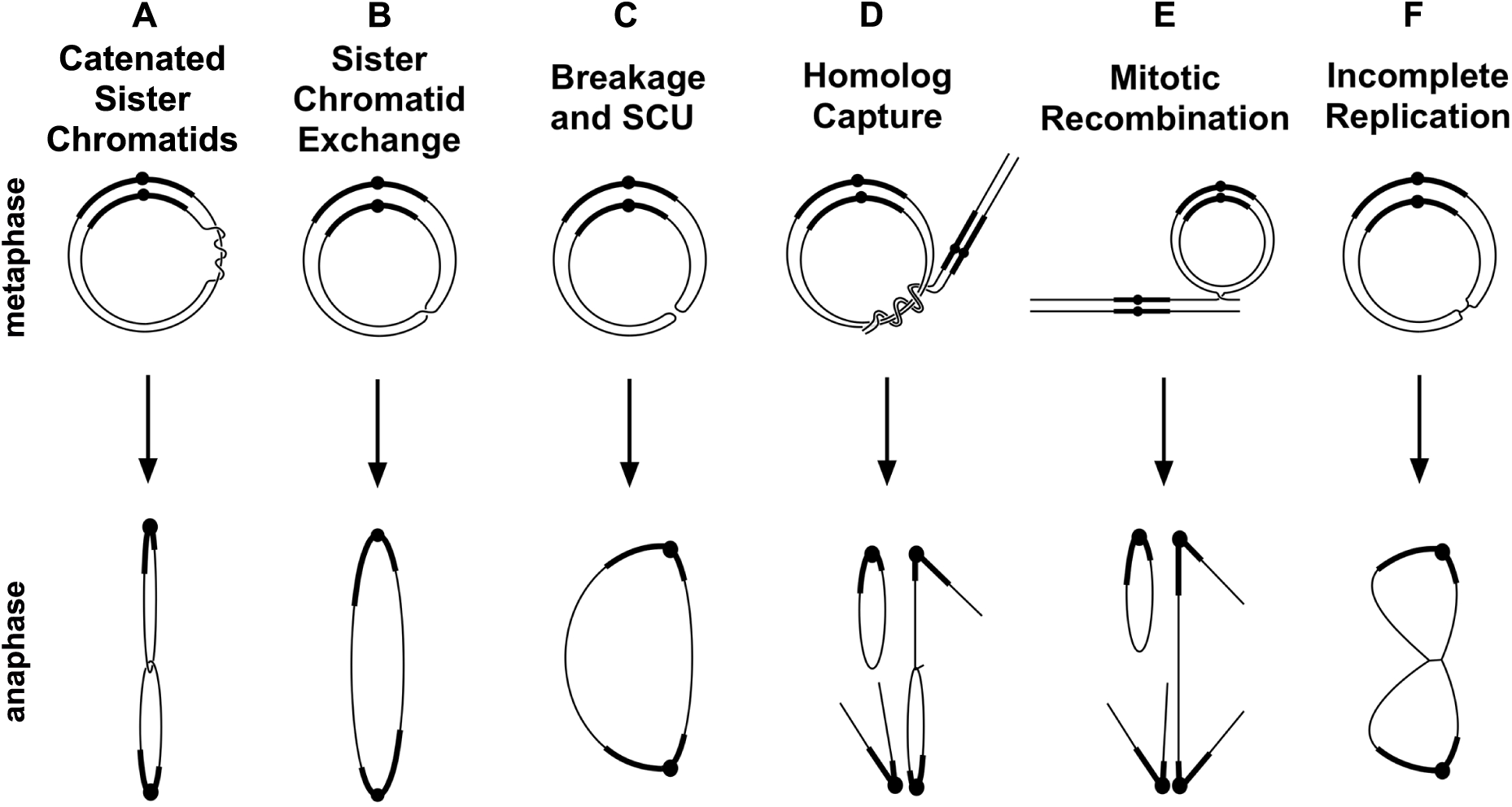
Mitotic misbehaviors of ring chromosomes. Six events that can produce anaphase chromosome bridges are shown.

In *Drosophila*, sex mosaics (gynandromorphs) are sometimes produced by early mitotic loss of ring-*X* chromosomes (Morgan 1926; Griffen and Lindsley 1946; Hinton 1955). Partly because of this interesting and experimentally useful property, they have been intensely studied (Hall et al. 1976; Leigh 1976; Zalokar et al. 1980). Ring-*X* chromosomes in *Drosophila melanogaster* show frequent anaphase chromatin bridges in embryos and nuclear fallout from the cortex during the late syncytial divisions, leaving gaps in the nuclear array at blastoderm (Ferree et al. 2014). Ring autosomes have received far less attention. A complete ring-*3* chromosome was generated by Craymer, but it has not been studied, apart from noting that it too causes incomplete dominant lethality (Craymer 1984). Partial autosomal ring chromosomes, consisting of portions of chromosomes *2* or *3*, showed varying degrees of instability. This was detected either as mosaicism for a dominant marker gene, suggesting chromosome loss, as varying numbers of the chromosome in different cells of the same individual, or as bits of chromatin lying between poles at anaphase suggestive of a broken bridge (Hilliker 1978; Novitski and Puro 1978). Detailed observations of ring-X and autosome behavior would be useful to identify common, and possibly divergent, behaviors.

We undertook cytological analysis of ring-X*Y*, ring-2 and ring-3 chromosomes in *Drosophila melanogaster* (*R(1;Y)*, *R(2)* and *R(3)* respectively) to more fully identify and quantitate their mitotic behaviors. We observed instances of all three problems mentioned above (Fig. 1A-C) and identified sister chromatid catenation as the most frequent of the three. We also observed interactions between the ring and its homolog (Fig. 1D, E), including a surprisingly common event that we call homolog capture leading to catenated homologs, which is most likely a consequence of mitotic pairing.

## Materials and Methods

*Ring chromosome construction.* Construction of the *R(1;Y)* chromosomes has been described (Golic and Golic 2010). Briefly, I-*Cre*I meganuclease was used to induce exchange between an attached-*XY* chromosome and *In(1)EN* to generate a tandem metacentric attached-*X* bearing a large portion of the *Y.* Meiotic recombination then generated single ring-X*Y* chromosomes. The two versions used in this work, *R(1;Y)6AX2* and *R(1;Y)11AX2* are X-rayed derivatives, generated because the original chromosomes were extremely lethal (Hill and Golic 2015). The changes responsible for decreased lethality have not been fully characterized, apart from *6AX2* having a significant deletion of heterochromatin.

*R(2)* and *R(3)* chromosomes were constructed by using FLP recombinase to catalyze exchange between *FRT-*bearing *RS5r* and *RS3r* constructs located near the left and right tips of each chromosome (Golic and Golic 1996; Ryder et al. 2007). FLP was expressed using a *ß2t-*promoted FLP construct, which gives post-meiotic expression in the male germline (Golic et al. 1997). For *R(2)*, the elements *P{RS5}5-HA-1693* at cytological locus 21B4 and *P{RS3}CB-6716-3* at 60F5 were used. These were obtained from the Kyoto, Japan, stock center (https://kyotofly.kit.jp/cgi-bin/stocks/index.cgi) as stocks 125348 and 124193, respectively. For *R(3)*, *P{RS3}CB-5511-3* at 61B1 and *P{RS5}5-HA-1486* at 100E1 were used. Kyoto stock numbers were 123669 and 125206, respectively. Ring chromosomes were recognized as *white^+^* recombinants and confirmed cytologically.

*R(1)2* was described by Schultz and Catcheside (Schultz and Catcheside 1937). *R(3)C* refers to the ring-*3* constructed by Craymer (1984). Both were obtained from the Bloomington, IN, Drosophila Stock Center.

*Cytological methods.* For whole embryo cytology, embryo fixation was performed according to Embryo Fixation Method 3 in Protocol 9.3 from Rothwell and Sullivan (Rothwell and Sullivan 2000), except that between Steps 1 and 2, embryos were incubated in a solution of heptane saturated with 37% formaldehyde for 5’ to prevent the occurrence of artifactual chromatin bridges. Embryos were stained with DAPI in 1X PBS and examined on an Olympus IX81 microscope equipped with a DSU spinning disc system at multiple magnifications.

Live analysis of embryonic mitoses with R(3) was accomplished by crossing *R(3)/TM6* males to females carrying an *H2Av-GFP* transgene. Mothers were allowed to lay eggs for 30 minutes on a small petri dish containing food. The petri dish was changed every 30 minutes. After 4 repetitions freshly laid eggs were collected. The eggs were dechorionated manually on double-stick tape. Electrical tape was placed on top of a glass coverslip and a small rectangle was cut out. A thin layer of heptane glue was applied within the rectangle and the dechorionated embryos were placed on the dried glue. A small amount of halocarbon oil was placed on top of the embryos to prevent desiccation and covered with a small piece of oxygen-permeable membrane. Gentle pressure was applied to ensure the embryos were stuck to the glass coverslip. The embryos were examined on the Olympus IX81/DSU.

For high resolution cytology of embryonic mitoses, the squash protocol of Gao *et al*. (Gao et al. 2009)was followed except that colchicine and hypotonic treatments were eliminated. Slides were mounted with Vectashield containing 5 µg/ml DAPI and examined with a Zeiss Axioplan microscope with a 100X/1.3 Plan Neofluar objective.

Cytology of mitoses in larval brains was carried out following the protocol of Gatti and Pimpinelli (Gatti and Pimpinelli 1983). For metaphase preparations, larvae were dissected in either 0.7% NaCl or 1X PBS; brains were transferred to 0.5% NaCitrate for 10’, fixed briefly in an 11:11:2 solution of acetic acid:methanol:water, then squashed in a drop of 45% acetic acid under a siliconized cover slip. The slide was frozen on dry ice, the cover slip popped off with a razor blade and the slide allowed to air dry. Slides were mounted with Vectashield containing 5 µg/ml DAPI and examined with a Zeiss Axioplan or Axio Observer microscope with a 100X/1.3 Plan Neofluar objective. For anaphase preparations, after dissection, the 10’ NaCitrate incubation was eliminated and replaced with a brief, few seconds, wash in distilled water, followed immediately by the fixation in the 11:11:2 solution and subsequent steps.

## RESULTS

For this work we generated ring versions of each major chromosome in *Drosophila melanogaster* (Fig. 2). *R(1;Y)11AX2* and *R(1;Y)6AX2* were constructed as previously described. *R(2)* and *R(3)* were constructed using FLP recombinase and *FRT*s located near the tips of the chromosomes. *R(3)C* is the ring chromosome constructed by Craymer. They all exhibit strong dominant lethality which is primarily embryonic (Hinton 1955; Stone 1982; Ferree et al. 2014). *R(1)2* was discovered nearly a century ago and has been maintained in stock since. It shows virtually no zygotic lethality.

**Figure 2:**
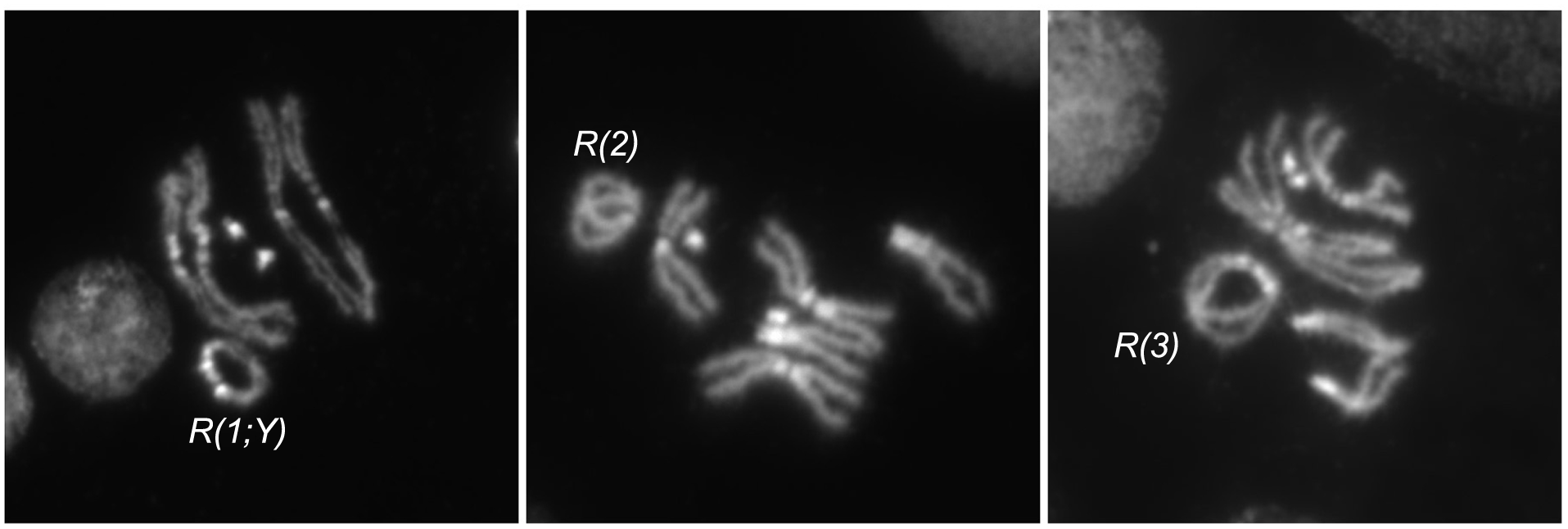
Cytology of ring chromosomes. Metaphase figures of three ring chromosomes used in this work.

### Ring chromosome mitotic defects in embryos

To identify the array of possible mitotic defects that produce anaphase chromosome bridges we examined embryos from crosses of *R(1;Y)11AX2* males to *y w* (control) females (Table 1). Similar to previous findings (Hinton 1959, Ferree et al. 2014), only 3% of embryos showed abnormalities in the first 4 mitotic divisions. However, beginning around the fifth division, 36% of the embryos from this cross exhibited problems, with missing nuclei and frequent chromosome bridges at anaphase. Several different bridge phenotypes were apparent, including thick bridges, thin bridges and double bridges (Fig. 3A). Since the *Y* portion of the *R(1;Y)* chromosome stains brightly with DAPI we were also able to identify instances of nondisjunction, where both copies went to the same daughter nucleus (gain in Fig. 3A), and chromosome loss, where the ring chromosomes did not make it to either daughter nucleus (loss in Fig. 3A). There were also frequent large gaps in the nuclear array (*aka* nuclear fallout) as embryos approached blastoderm (Fig. 3B).

**Figure 3:**
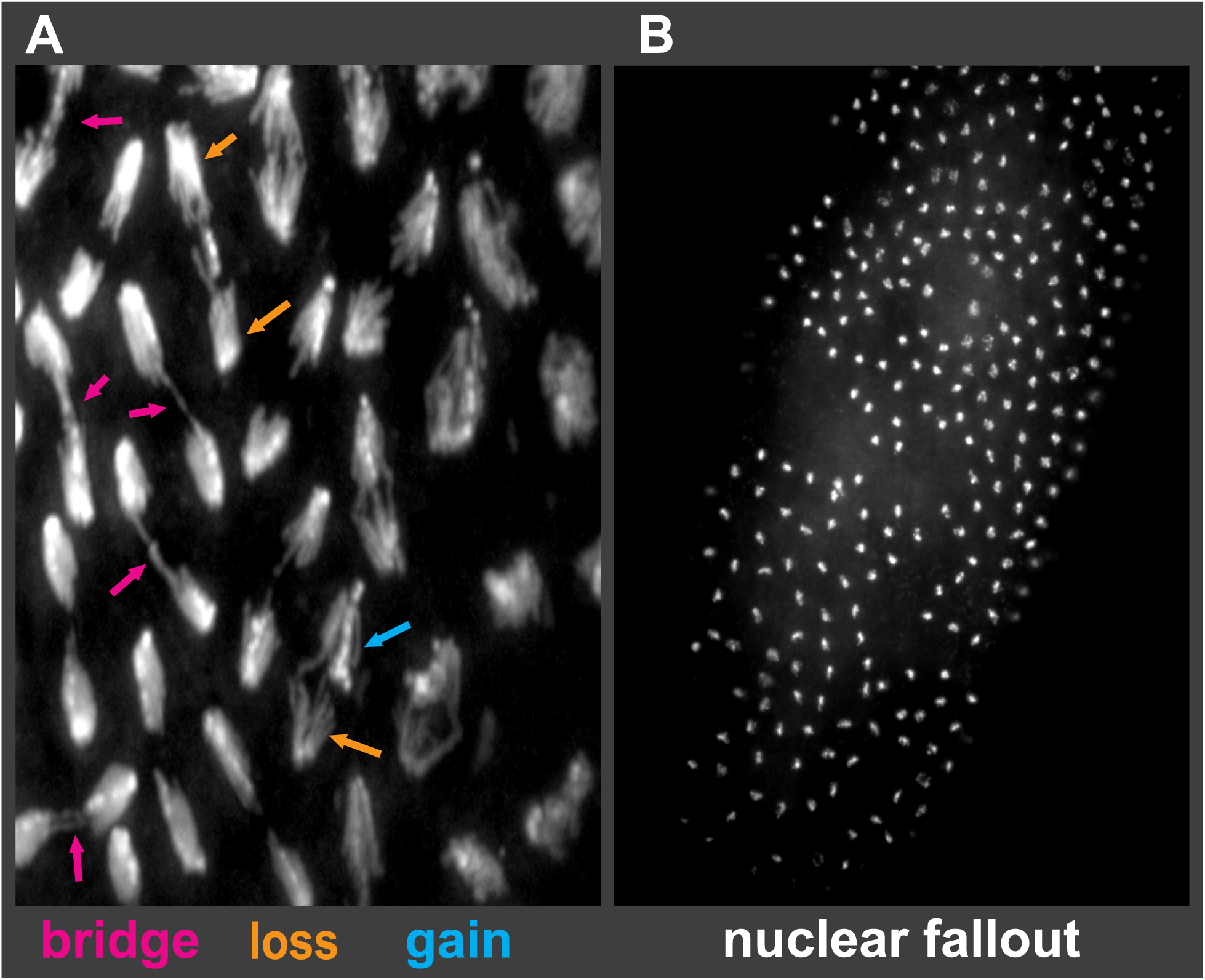
Embryonic mitosis phenotypes of *R(1;Y)* embryos. (A) Embryonic mitoses showing anaphase bridges of varying types and chromosome loss and gain events. (B) Typical blastoderm embryo showing large regions lacking nuclei owing to nuclear fallout as a response to unresolved bridges.

**Table 1:**
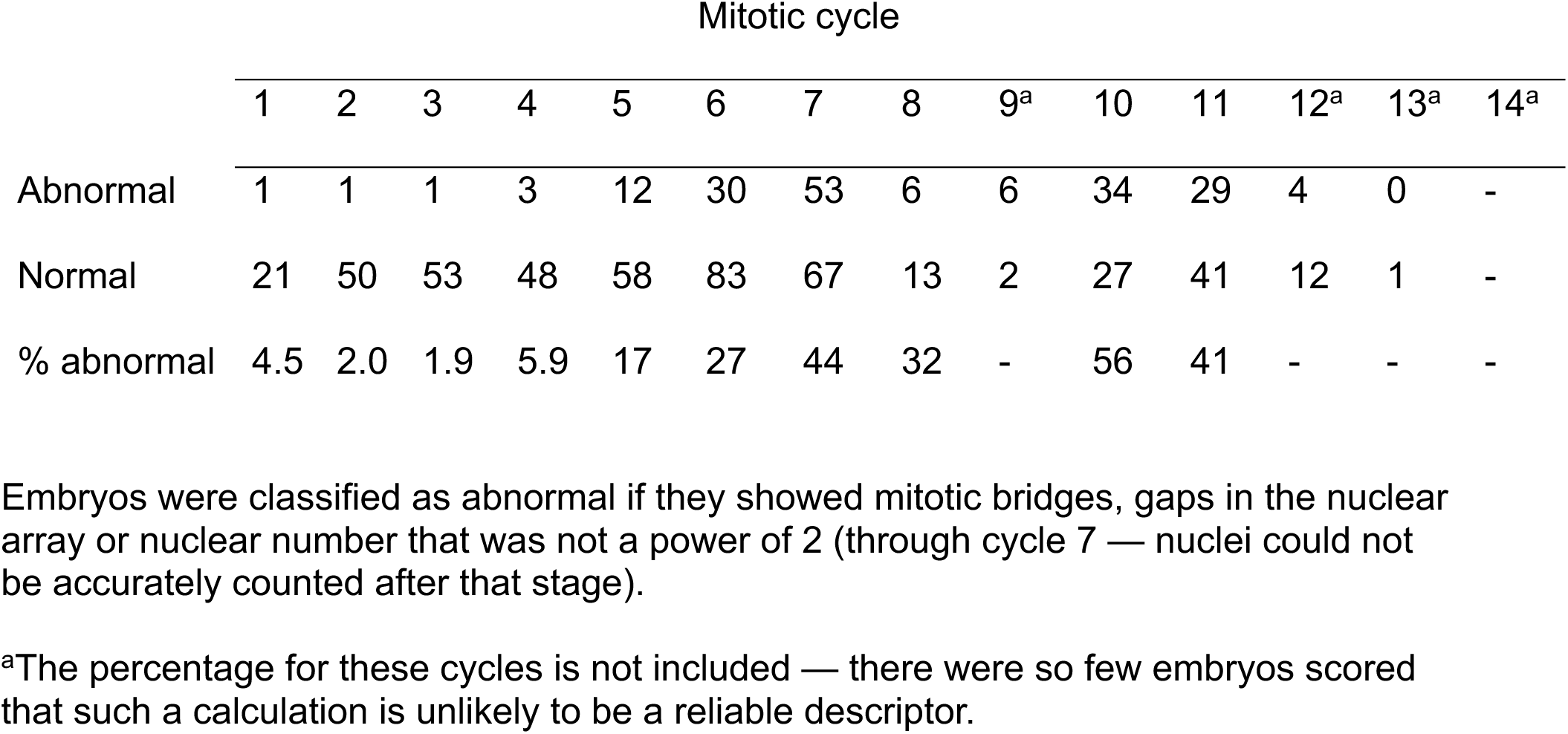
Abnormal embryos from R(1;Y)11AX2 males x y w females by mitotic cycle.

In contrast, *R(1)2* embryos showed very few defects, though occasional bridges were seen. Embryos from *y w/Y* control fathers were essentially normal, with only occasional nuclei showing any abnormalities. Approximately 5% of embryos from *y w* fathers showed no evidence of nuclear division and were likely unfertilized eggs.

Mitotic chromosome bridges in *R(1;Y)11AX2/y w* embryos were examined more closely by squashing embryos and staining with DAPI. Half of the mitoses from ring-bearing embryos showed some variety of chromatin bridge (60/121). Although the majority were not clearly classifiable, in some cases chromatid strands within the bridges could be resolved making it possible to deduce the likely cause of the bridge. Of those that could be classified, most (15/18) appeared to result from catenated, or interlocked, rings (Fig. 1A; Fig. 4A,B). There were also bridges with a thin connection between daughter chromatids (Fig. 4E, arrow) which may represent stretched or resolving catenanes. Double bridges were clearly identified in only a small number of cases (3/18). Those produced by SCE can be recognized because they have two equal-length bridges oriented in opposite directions (Fig. 1B). In the *R(1;Y)11AX2* chromosome, where the DAPI-stained heterochromatin is arranged asymmetrically about the centromere, these bridges have a unique appearance, and a mitotic figure matching this expectation was observed (Fig. 4C). Chromosome breakage and sister chromatid union (SCU; Fig. 1C) will also produce double bridges, but these bridges will not show the reversed symmetry of bridges created by SCE, and there was also an example of this (Fig. 4D). The third anaphase with a double bridge could not be distinguished.

**Figure 4:**
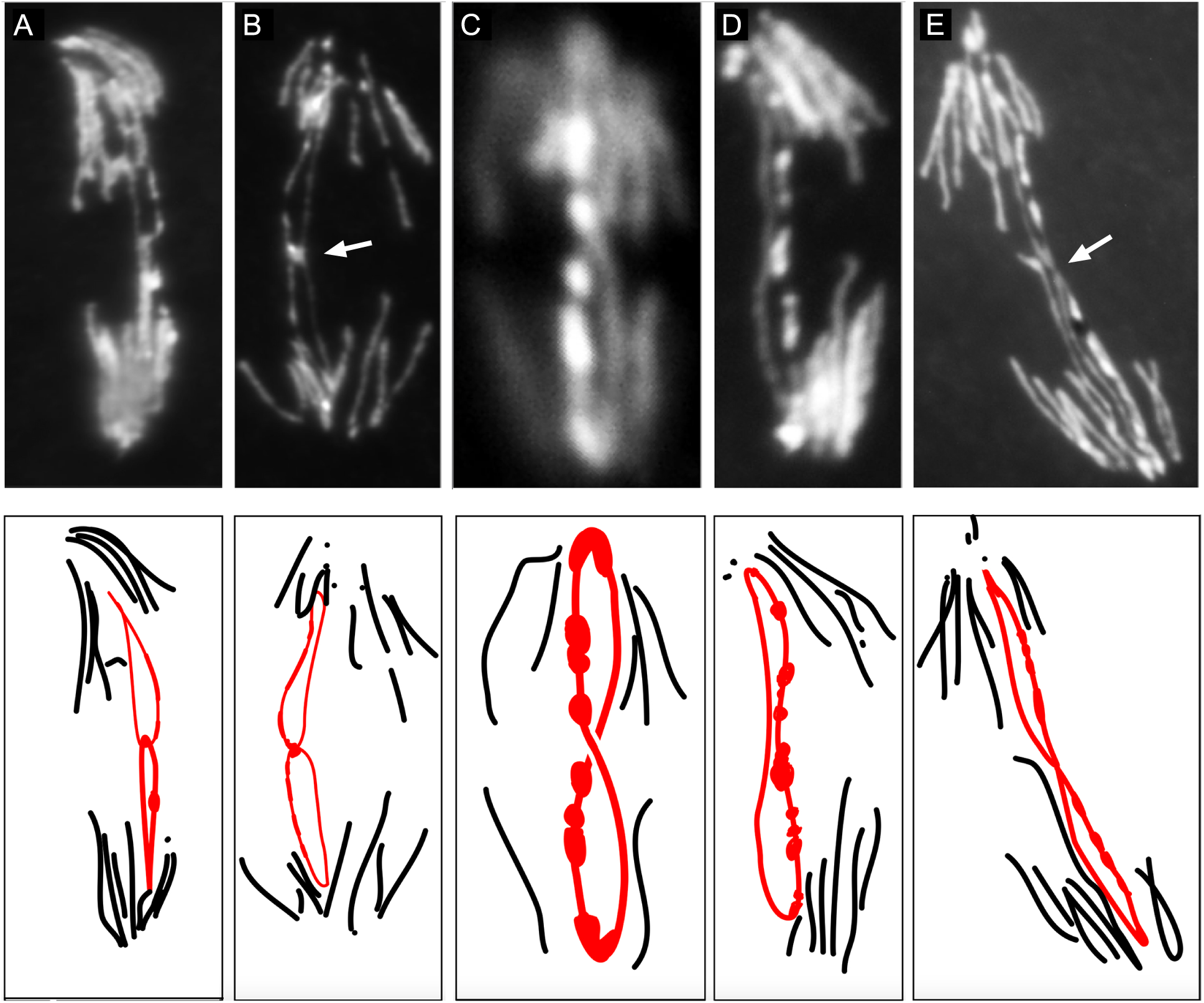
*R(1;Y)* embryonic mitoses from a squash preparation showing different types of anaphase bridges. Below each photograph is a schematic interpretation with the ring chromosome anaphase bridges in red. Shown are: (A,B) sister chromatid catenanes (arrow in B indicates bright region at junction); (C) a double bridge generated by SCE; (D) A double bridge generated by SCU; (E) sister chromatids with an attenuated connection.

We also examined fixed embryos from crosses of *R(3)/TM6* males x *y w* females and, similar to *R(1;Y),* saw numerous chromosome bridges and large gaps in the nuclear array in many of the embryos, along with many normal embryos, which we presume to be those that received *TM6*.

Since fixed images do not directly show the fates of chromosome bridges we captured live time-lapse images of mitoses in pre-blastoderm *R(3)* embryos (Fig. 5). The large size of the *R(3)* chromosome makes it relatively easy to visualize during mitosis. A few nuclei did not divide while under observation, but the majority (66%) of dividing nuclei did form anaphase bridges during mitosis, and all but one of these were clearly a result of sister chromatid catenation (Table 2). In Fig. 5A,B the analogy to links in a chain is striking: during anaphase the connected rings are oriented orthogonally, just as they would be in a stretched chain. Most catenanes (82%) were resolved during anaphase (Fig. 5A), allowing daughter nuclei to separate. Bajer observed catenated ring sister chromatids resolving in a similar fashion in cells of the Blood Lily (Bajer 1963).

**Figure 5:**
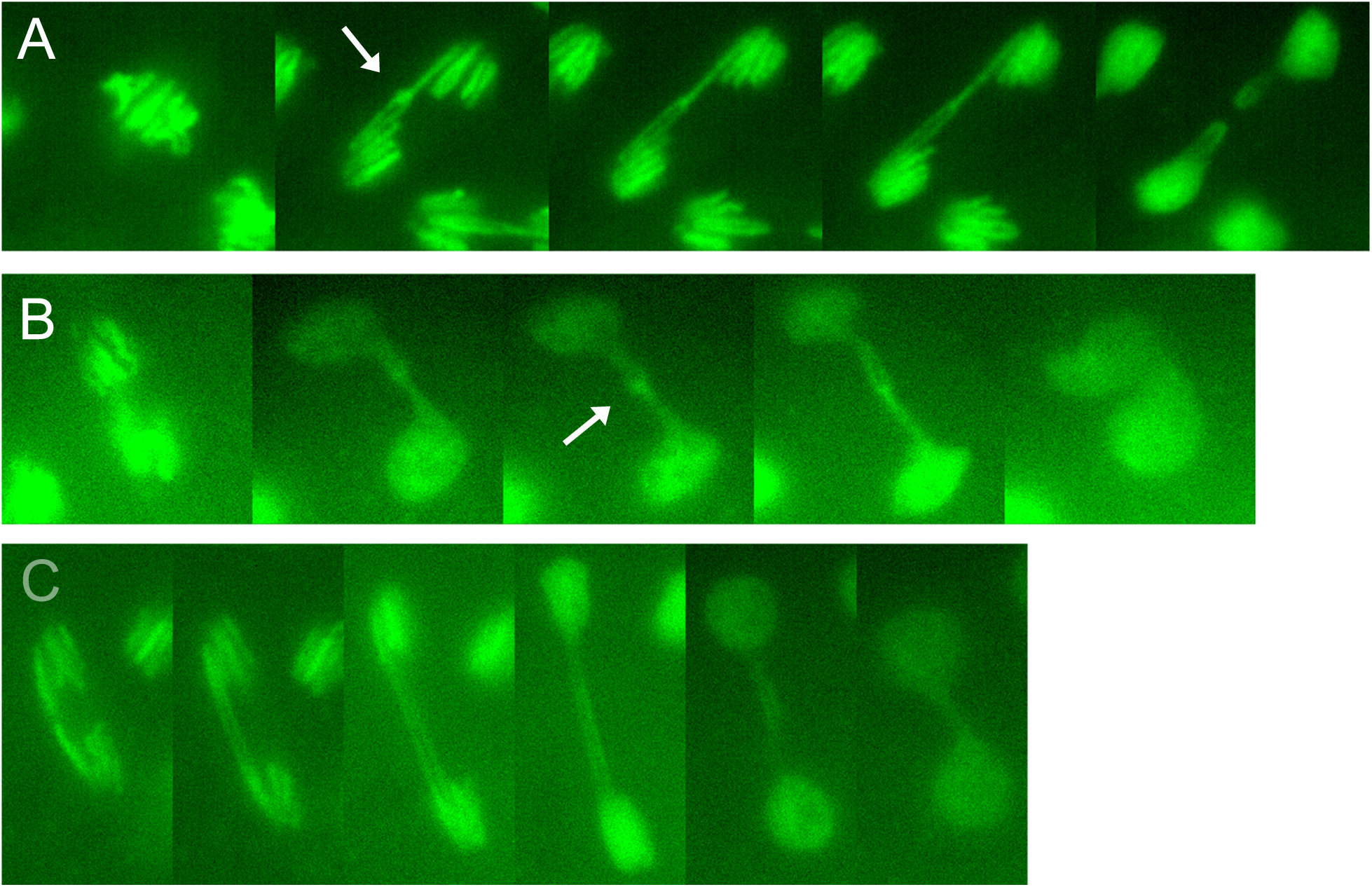
*R(3)* embryonic mitoses from time-lapse video, showing: (A) catenated sister chromatids that stretch and then resolve into separate and intact daughter rings; (B) catenated sister chromatids that do not resolve followed by daughter nuclei moving back together; (C) a dicentric double bridge, likely formed by SCE or SCU.The arrows in A, B indicate regions of concentrated chromatin at the junctions of sister chromatids.

**Table 2:** Live analysis of mitoses in *R(3)* embryos.

| total nuclei | normal | resolved catenane | unresolved catenane | double bridge | failure to divide |
| --- | --- | --- | --- | --- | --- |
| 94 | 19 (20%) | 50 (53%) | 11 (12%) | 1 (1%) | 13 (14%) |

A significant minority (18%) of catenated sisters did not resolve, and unseparated daughter nuclei formed a dumbbell shape after mitosis (Fig. 5B). Only one dividing nucleus exhibited a parallel double bridge (Fig. 5C), indicating that SCE is an infrequent contributor to bridge formation in embryos with *R(3)*.

### Mitotic defects in larval brains

When larval mitoses with ring-X chromosomes were previously examined by others, anaphase chromatin bridges were seen at rates of 3-22% — significantly less frequent than we observed in embryos (Braver and Blount 1950; Hinton 1955; Brosseau 1966). Although the chromosomes previously examined are not directly comparable to those we constructed, those earlier observations raise the possibility that embryonic mitoses are more frequently affected by ring misbehavior. Since the majority of ring-dependent zygotic death occurs in embryos it seemed possible that mitotic problems would be ameliorated in later stages of development. One potential explanation is that embryonic divisions are extremely rapid, with cycle times as short as five minutes, while larval mitoses are on the order of several hours. The longer mitotic cycles in larval brains might allow cells to more reliably disentangle sister chromatids during mitosis. To test this, we dissected brains from ring-bearing third instar larvae and examined anaphase figures.

In brains of *R(1;Y)6AX2* and *R(1;Y)11AX2* larvae nearly half of the anaphase figures were abnormal (Table 3), showing some form of chromatin bridge between daughter nuclei, similar to the frequency of bridges seen in embryos. Some of these were obviously catenated rings while others could be classified as clear double bridges resulting from either SCE or SCU, but most were not clearly classifiable. As was the case in embryos, mitoses showing anaphase bridges were much less frequent with *R(1)2* but were still seen in a few percent of mitoses.

**Table 3:** Anaphase figures in larval brains.

| ring | N | Anaphase phenotypes |  |  |  |  |  |  | unscorable<sup>b</sup> |
| --- | --- | --- | --- | --- | --- | --- | --- | --- | --- |
|  |  | normal | double<br>bridge | catenane | homolog<br>capture<sup>a</sup> | other<br>bridge | broken<br>bridge | total<br>bridges |  |
| *R(1)2* | 13 | 98 | 0 | 1 | 0 | 7 | 0 | 8% | 18 |
| *R(1;Y)6AX2* | 6 | 27 | 3 | 0 | 0 | 18 | 1 | 45% | 4 |
| *R(1;Y)11AX2* | 11 | 26 | 3 | 1 | 0 | 13 | 5 | 46% | 8 |
| *R(3)* | 4 | 50 | 9 | 11 | 7 | 32 | 2 | 55% | 26 |

N, number of brains scored
<sup>a</sup> For ring-*X* or *XY* chromosomes, homolog capture might be influenced by whether the homolog is an *X* or *Y*. In the majority of cases, brains of female larvae were scored (11/13 for *R(1)2*, 5/6 for *R(1;Y)6AX2*, 6/11 for *R(1;Y)11AX2*). However, definitive examples of homolog capture were only observed with *R(3)*.
<sup>b</sup> Most of these were early anaphase, where the chromosomes had not separated sufficiently to determine whether there was a bridge.

With *R(3)*, normal anaphase figures (Fig. 6A) were in the minority (Table 3) — more than 50% of anaphase figures showed bridges. We only quantitatively scored the *R(3)* that was constructed in our laboratory, but similar mitotic problems were seen during our examination of *R(3)C*. In some anaphases, sister rings appeared to be catenated (Fig. 6B). Although catenanes could potentially be resolved by topisomerase II, either before or during mitosis, with *R(3)* they accounted for a little more than half of the scorable anaphase bridges. Sister chromatid catenanes sometimes persisted into telophase or beyond (Figure 6C). McClintock similarly saw occasional cases where bridges persisted into the following interphase in corn (1938). There were bridges between *R(3)* sister chromatids with a thin connection in the middle (Fig. 6D), similar to those we saw in *R(1;Y)* embryos. Connected rings also caused occasional nondisjunction (Figure 6E), leading to rare trisomy (Figure 6F).

**Figure 6:**
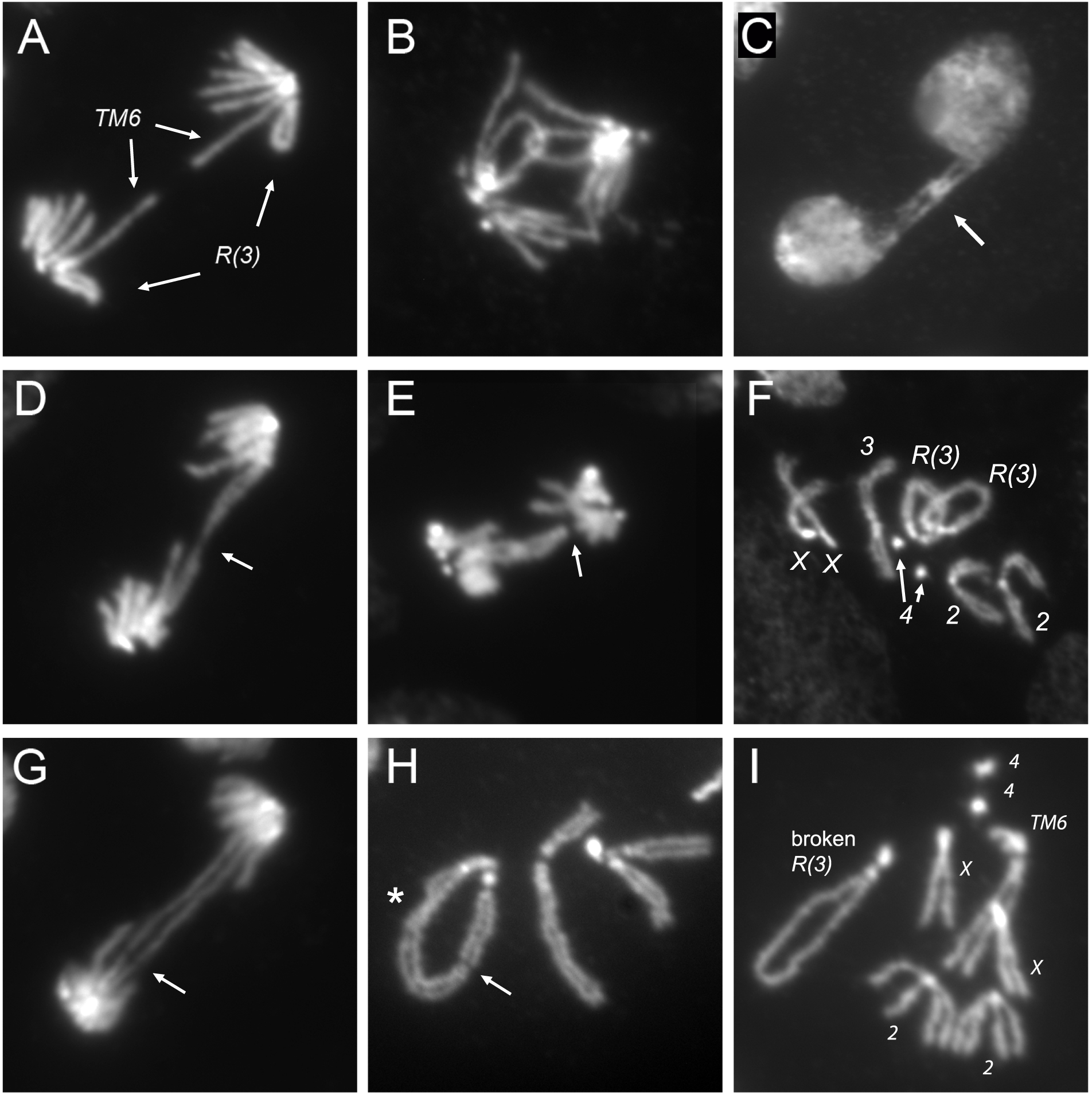
*R(3)* larval mitoses. Figures are from *R(3)/TM6* or *R(3)/+*. Shown are: (A) normal anaphase; (B) catenated sister chromatids in anaphase; (C) catenated sister chromatids (arrow) connecting interphase daughter nuclei that have progressed to the subsequent interphase; (D) sister chromatids connected by an attenuated thread of chromatin (arrow); (E) catenated sister chromatids with one sister having apparently lost connection to the spindle (arrow); (F) *R(3)/R(3)/+* trisomy; (G) a double dicentric bridge, likely resulting from SCE, with incipient break (arrow); (H) a break in *R(3)* (arrow) with sisters fused on one end of the break; also indicated (asterisk) is a site where sister chromatids are twisted about each other; (I) another example of a broken *R(3)* which, in this case, was recovered as a spontaneous opening of the ring chromosome through the germline.

In some cases a double bridge was clearly visible (Fig. 6G). It seems probable that at least some were produced by SCE since they often showed equal length bridges. This example also appears to show an incipient break (Fig. 6G, arrow) and we did see occasional examples of broken chromosomes in metaphase nuclei. For example, Fig. 6H shows *R(3)* with a break at a position similar to that illustrated in Fig. 1C, but with SCU on only one side of the break. Fig. 6I shows an instance of a spontaneously opened ring where the break occurred in heterochromatin near the heterochromatin/euchromatin junction. As with sister chromatid catenanes, there were also a few examples of unbroken double bridges persisting to interphase (not shown), indicating that breakage may not be inevitable.

### Ring chromosome interactions with the homolog generate anaphase bridges

In metaphase nuclei there were many examples where the ring chromosome appeared to encircle its linear homolog (Fig. 7A-C). This occurred with all ring stocks examined: *R(1;Y)/FM7, R(2)/CyO* and *R(3)/TM6* (Table 4).

**Figure 7:**
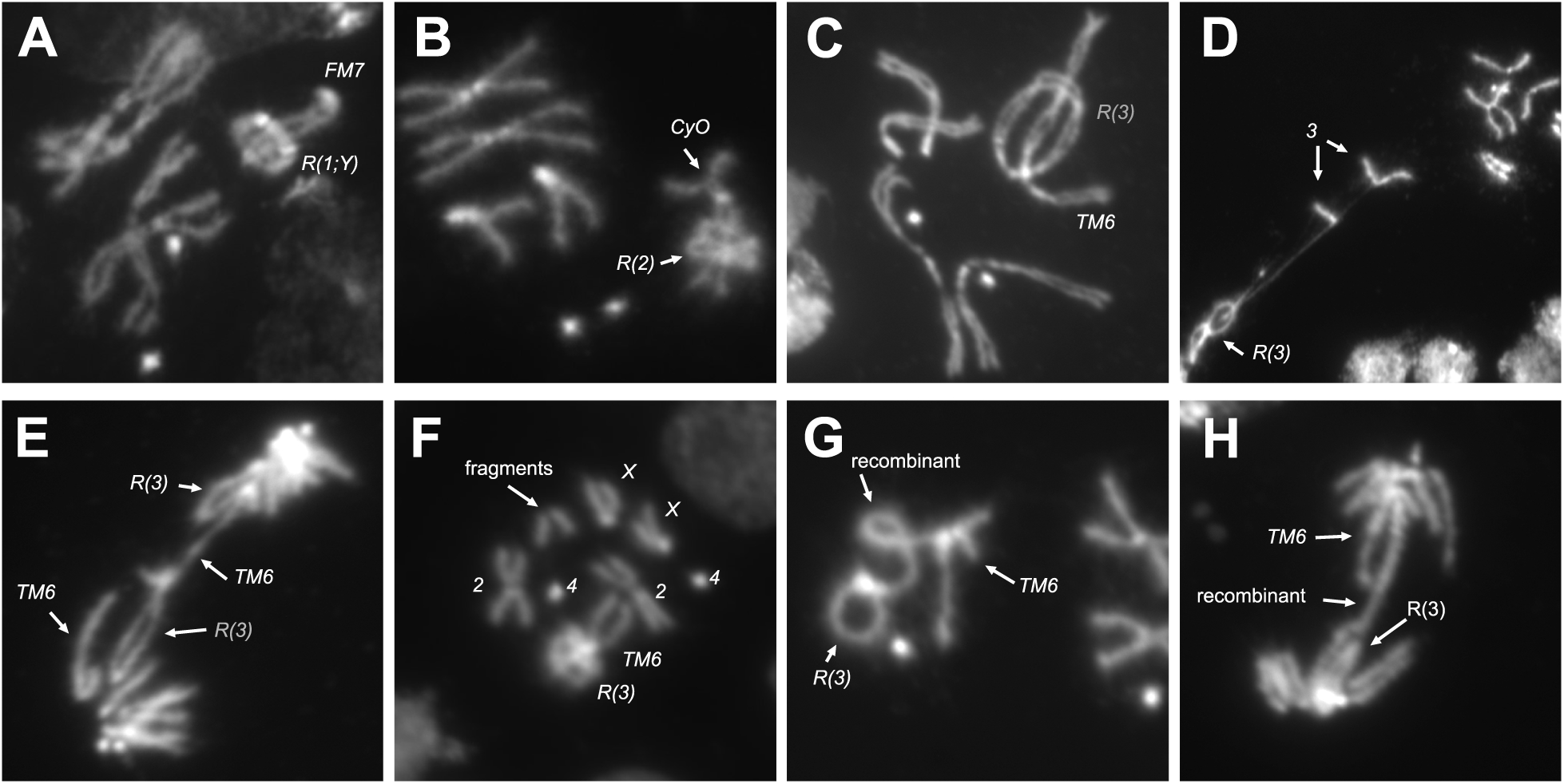
Interactions of ring chromosomes with their homologs. Ring chromosomes are frequently wrapped around their linear homologs in metaphase, with examples of: (A) *R(1;Y)/FM7;* (B) *R(2)/CyO*; (C) *R(3)/TM6.* Homolog capture by *R(3)* in: (D) *R(3)/+*; (E) *R(3)/TM6*. (F) Metaphase showing extra chromosome fragments. Mitotic recombination events between *R(3)* and the *TM6* homolog at: (G) metaphase; (H) anaphase.

**Table 4:** Encirclement of linear homolog by ring chromosomes.

|  | metaphases scored | encircled |
| --- | --- | --- |
| *R(1;Y)/FM7* | 16 | 38% |
| *R(2)CyO* | 131 | 41% |
| *R(3)/TM6* | 81 | 41% |

Although it seems that the linear chromosome should easily slide out of the ring when pulled at anaphase, this may not always be so. In one nucleus, where the chromosomes were stretched and separated by squashing, the ring chromosome pulled one sister of its linear homolog with it, leading to the conclusion that the ring and linear homolog were entangled or attached (Figure 7D). This “capture” of the linear homolog by the ring is not simply an artifact of squashing — there were several anaphase figures in which *R(3)* appears to have captured its linear homolog, creating an anaphase chromatin bridge (Figure 7E, Table 2). There were a few metaphase or anaphase nuclei with additional acentric chromosome fragments (Fig. 7F). We did not determine the identity of these fragments, but they might represent the terminal region of the linear chromosome *3* homolog that was torn away from a sister nucleus after capture by the ring chromosome during the previous mitosis.

Finally, there were instances of recombination between the ring and its homolog, which generates a single chromatin bridge in anaphase (Figure 7G, H). Recombination between homologs was rare — we saw five instances among thousands of metaphase and anaphase nuclei: the two shown in Fig. 7 in *R(3)/TM6* flies and three others in *R(1;Y)* homozygotes, producing double-sized ring dicentric chromosomes (not shown).

### Catenation of homologous chromosomes as a consequence of mitotic pairing

Homolog capture events, such as seen in Fig. 7E, could have multiple origins. For instance, recombination intermediates between homologs that are not resolved prior to anaphase might produce such bridges. This seems unlikely, primarily because of the rarity of mitotic recombination relative to instances of homolog capture. A more probable source of these connections is the homolog pairing that occurs normally in mitotic cells. In this intimate pairing the linear homolog might twist around its circular partner, and when pairing is relaxed in mitosis the linear chromosome may end up in the middle of the ring (Fig. 8A). Anaphase entanglements might occur if the linear chromosome wrapped its homolog with multiple twists that were not resolved before anaphase (Fig. 8B). In support of the idea that this reflects homolog pairing, examples of the ring surrounding a chromosome other than its homolog were extremely rare, with only five examples seen in thousands of metaphases.

**Figure 8:**
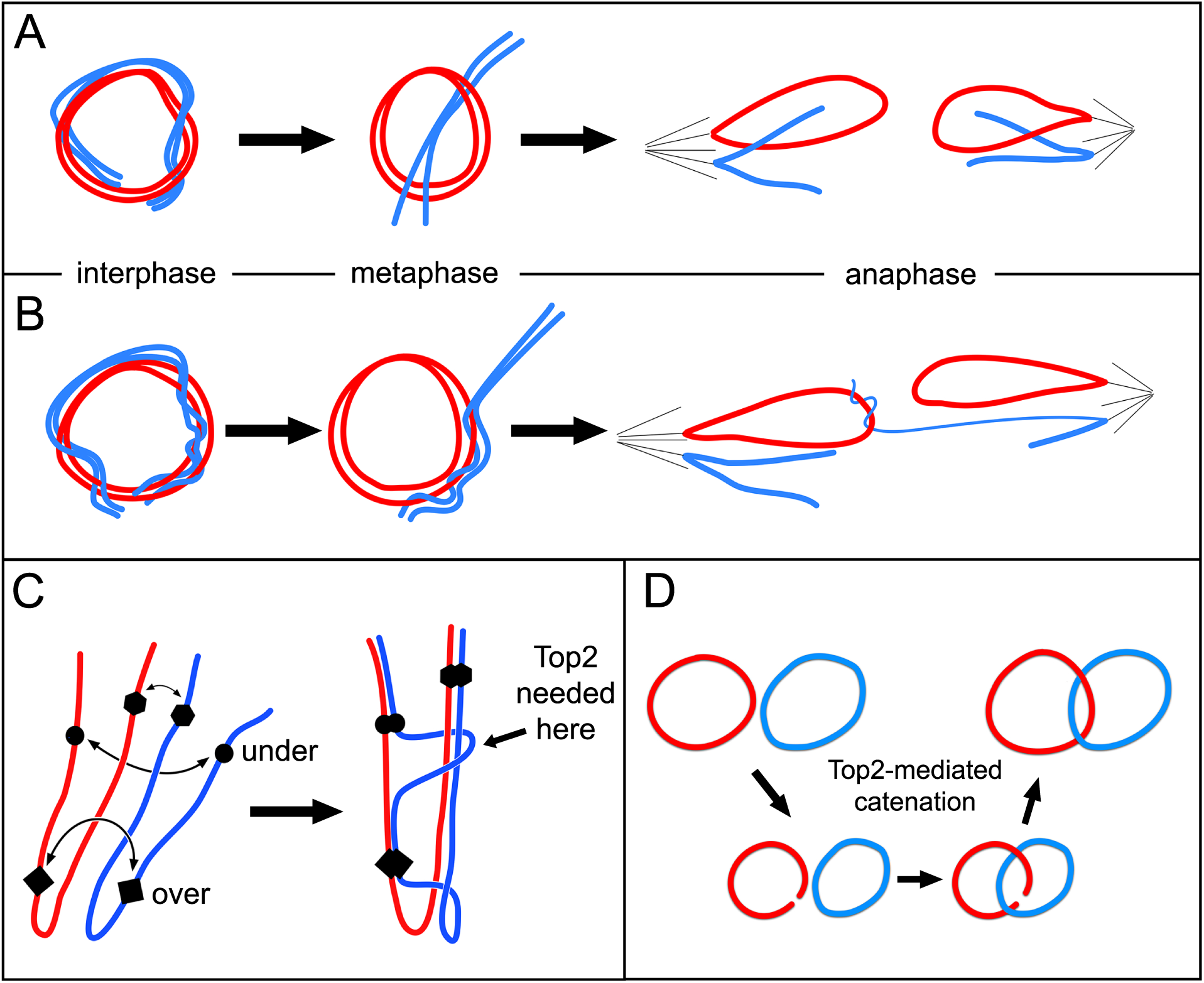
Chromosome pairing drives entanglement and catenation of homologs. Red and blue are used to indicate homologous chromosomes. (A) Pairing of ring and linear homologs may end with ring encircling linear homolog at metaphase. (B) Extensive wrapping of linear about ring chromosome may lead to capture of the linear chromosome by the ring to generate an anaphase chromosome bridge. (C) Three hypothetical pairing sites are indicated on homologous chromosomes lying in a Rabl configuration after mitosis. Forces that drive pairing may produce tangles that require chromosomes to pass through each other to achieve full pairing, a process that may require topoisomerase II (Top2). (D) Topoisomerase II activity mediates catenation of ring homologs.

Cytologically, pairing in diploid mitotic cells is typically recognized as the relative proximity of condensed homologs at metaphase (*e.g.,* Fig. 9A; Stevens 1908). However, this is a mere shadow of the intimate pairing that occurs in interphase. This is readily seen with chromosomes in mitotic prophase, where homologs are paired so closely as to seem a single chromosome, except for centric heterochromatin which is not paired at this stage (Fig. 9B). The intimate pairing of homologs in interphase can also be seen by fluorescent *in situ* hybridization, where often a single dot of hybridization is seen even though two alleles are present (Hiraoka et al. 1993). If the forces of pairing that drive juxtaposition of homologous sequences push chromosomes against each other topoisomerase II may be needed to pass one chromosome across another to achieve full pairing (Fig. 8C). Multiple passages could produce homologs that are intertwined. In fact, prophase chromosomes often give the appearance of homologs twisted about each other (Fig. 9B, arrow). In some cases non-sister chromatids appear more closely paired and twisted together than sister chromatids (Fig. 9C, arrows). It follows that in a ring/rod karyotype the linear chromosome may find itself wrapped around its ring homolog.

**Figure 9:**
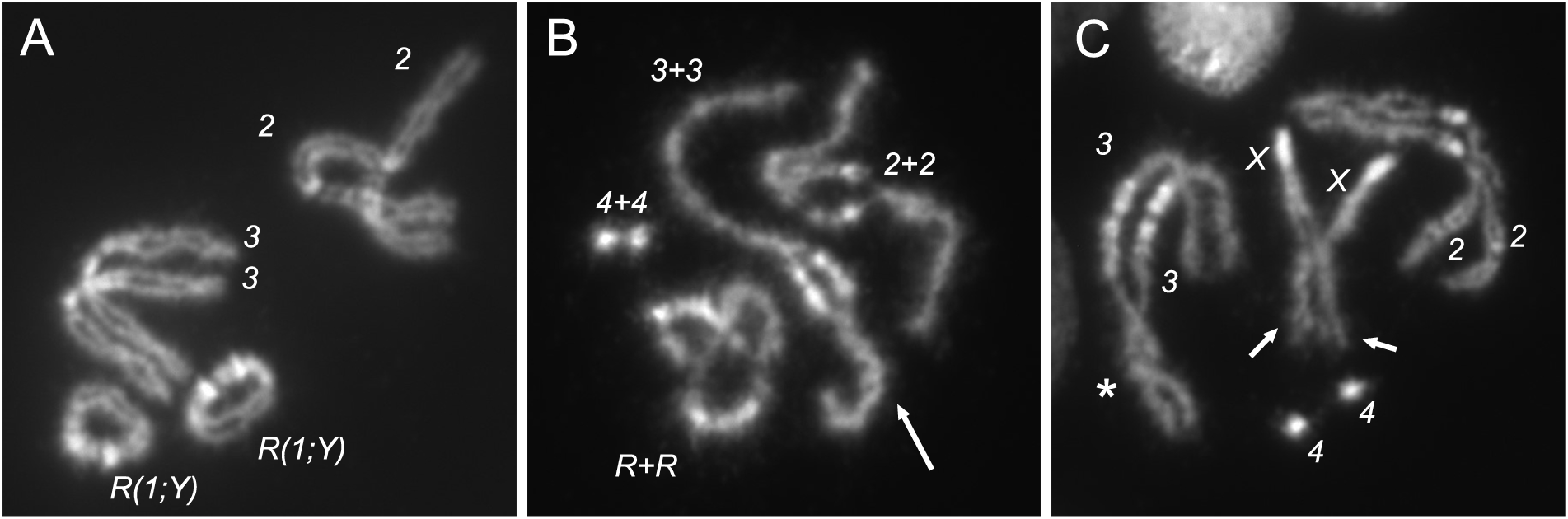
Mitotic pairing of homologs visualized at different stages of mitosis. (A) In metaphase squashes, chromosomes are often found lying closer to their homologs than to heterologous chromosomes. (B) In prophase homologs are intimately paired and often appear as a single chromosome, apart from centric heterochromatin (seen as areas of bright DAPI staining), which is not paired at this stage. At this stage, chromosomes appear tightly wound around each other (arrow). (C) As cells progress towards metaphase, and pairing is dissolved, occasionally non-sister chromatids are seen to be more tightly associated than sister chromatids (arrows). Also shown is an example of homologs loosely twisted about each other (asterisk). This image is a non-ring genotype but was chosen because it illustrates this type of association.

To test the underlying hypothesis, that homolog pairing drives chromosomes to pass through each other, we examined metaphase spreads from *R(1;Y)/R(1;Y)* homozygous females (neither *R(2)* nor *R(3)* is homozygous-viable). Ring homologs could only become catenated if at least one ring opened, the other ring passed through that opening, and the first ring resealed (Fig. 8D). We found that 21% (195/754) of metaphases showed catenation of one or both non-sister chromatids (Fig. 10A-C). Another 26% (195/754) showed ring homologs lying on top of each other, but could not be definitively scored as catenanes, though we expect that a significant fraction of these were indeed catenated. In 54% of metaphases (404/754) the rings were clearly separate. In anaphase figures there were cases of catenated homologs and catenated sister chromatids (Fig. 11A and B, respectively), though the latter configuration might also be produced by homolog catenation if linked non-sister chromatids segregated apart, rather than together.

**Figure 10:**
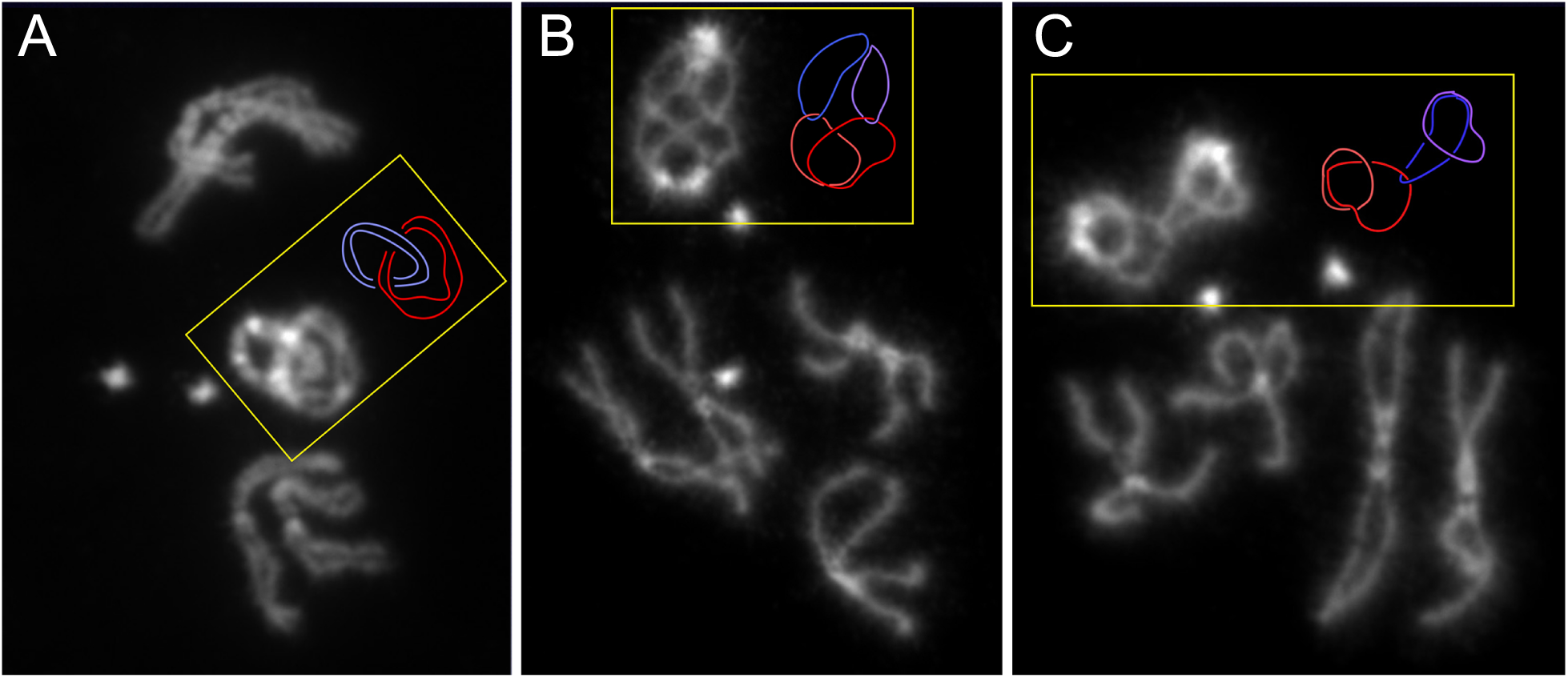
Ring chromosomes reveal catenation of homologs. *R(1;Y)11AX2/R(1;Y)11AX2* larval metaphases showing: (A) catenation of both sisters of both homologs; (B) catenation of each chromatid with one non-sister chromatid; (C) catenation of one chromatid with one non-sister chromatid. Shades or either red or blue are used to indicate sister chromatids in the accomanying schematic diagrams.

**Figure 11:**
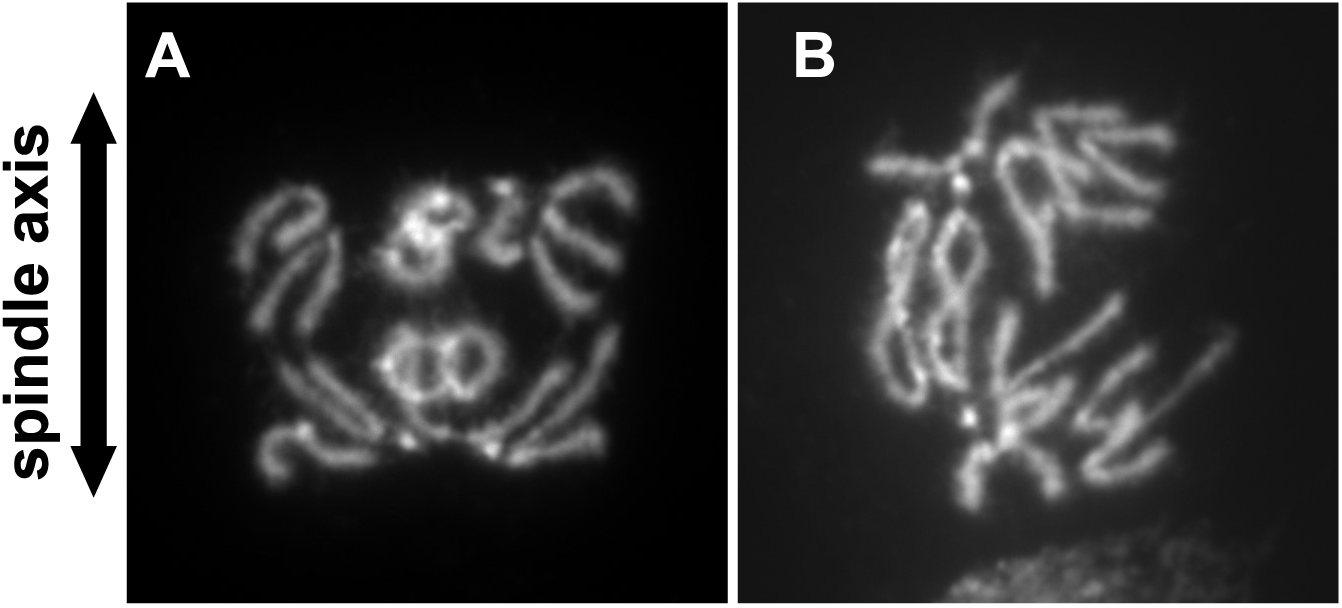
Anaphase figures from *R(1;Y)* homozygotes. Anaphase figures showing either: (A) catenation of non-sister chromatids; or (B) catenation of sister chromatids.

We conclude that mitotic pairing can drive frequent catenation of homologous chromosomes. The resultant intertwining or braiding of homologs is likely responsible for the homolog capture events we observed in ring/rod heterozygotes.

Unlike the brains of *R(3)/TM6* larvae, where chromosome *3* trisomy was extremely rare, triplo*-X* cells were seen in most brains of *R(1;Y)/R(1;Y)* homozygotes. Cells with this trisomy were found in 10/11 fully-scored *R/R* brains, and though always a small fraction, they frequently occurred as several trisomic cells in a single brain. In some cases these cells were adjacent in the squash preparation suggesting that they represented a small clone. Cells showing the complementary monosomy were comparatively infrequent, found in only 4/11 brains and always appeared as a single cell, confirming Hinton’s previous findings (1955). The cells showing *R(1;Y)* aneuploidy probably arose by nondisjunction relatively late in larval development, well after dosage compensation had been set in early embryogenesis. Late-arising trisomy and monosomy, but especially monosomy, are expected to be deleterious, accounting for the small clone sizes. We did, however, find brains that were entirely *XO* in 2/18 female larvae (as judged by the absence of testes), no doubt representing gynandromorph production from early loss of a *R(X;Y)* chromosome. It is possible that *R(3)* experiences equally frequent nondisjunction or loss, but the trisomic/monosomic cells are rarely seen because they are quite inviable.

## Discussion

It has been known for nearly a century that ring chromosomes produce chromatin bridges in mitotic anaphase (McClintock 1938). From extensive examination of ring chromosomes in corn, McClintock attributed the occurrence of anaphase bridges primarily to SCE (Fig. 1B), but also acknowledged that the cytological resolution could not exclude the possibility that some bridges were a result of catenated sister chromatids (Fig. 1A). SCE is normally quite rare in *Drosophila*, but, though still rare, does occur more frequently with ring chromosomes (Gatti et al. 1979). With ring-*X* chromosomes in Drosophila, double bridges consistent with production by SCE and catenated sisters have both been seen (Braver and Blount 1950; Brosseau 1966). On the other hand, Hinton saw no examples of catenated sisters, and proposed that chromosome breakage followed by SCU was the basis for ring-*X* chromosome misbehavior (Hinton 1955; Hinton 1957). More recently, an interaction between topoisomerase II and the 359 bp satellite of the Drosophila *X* chromosome was implicated in anaphase bridge formation (Ferree et al. 2014)

In our examinations of embryonic mitoses, bridges produced by sister chromatid catenanes were much more frequent than those produced by SCE though we also saw examples of the latter. Live analysis showed that more than 80% of catenanes that persisted to anaphase were resolved, almost certainly by the action of topoisomerase II, allowing separation of daughter nuclei. In larval brain squashes approximately 50% of anaphases showed chromatin bridges. Most were not clearly classifiable, but among those that could be classified there were approximately equal numbers of catenanes and dicentric double bridges.

Catenanes might arise simply by replication of a circular DNA molecule where the strands are wrapped around each other every 10 base pairs. Catenanes could also be produced by an even number of SCE events, where the even-numbered exchange resolves to generate topologically linked rings. We think it likely that most catenanes result from replication of the double helix, rather than SCEs. First, at least in embryos, double bridges were much less frequent than catenated sisters. Unless there exists some mechanism to enforce an even number of SCE events, the frequency of SCE seems insufficient to account for the high frequency of catenanes. Second, when catenanes were stretched, the junction between rings sometimes appeared much brighter than adjacent chromatin, suggesting multiple wraps of sister chromatids and an accumulation of knotted chromatin under tension (arrows in Fig. 4B, 5A,B), instead of the single loop that would be produced by two successive SCEs (as in Fig. 6B, for instance). Finally, if catenanes were the result of a single topological link of sister chromatids it is likely that topoisomerase II would easily resolve the catenated rings. That this was not always true suggests that catenanes could involve multiple wraps which are not easily resolved.

Double bridges produced by SCE or by SCU did occur, and were a larger fraction of anaphase bridges in larval mitoses than in embryos. It was not always possible to determine whether a bridge arose by SCE or SCU. For *R(1;Y)*, the large amount and asymmetric distribution of heterochromatin sometimes allowed them to be distinguished (as in Fig. 4). For *R(3)*, the two events could not necessarily be distinguished in anaphase, but SCU events were occasionally recognized in metaphase squashes (Fig. 6H). SCEs would be difficult to identify at metaphase by DAPI staining alone, as they might appear as a simple twist between sisters and would not be easily distinguished from catenated sisters. If the double bridges we saw in larval brains were a result of SCE, then the *R(1;Y)* and *R(3)* chromosomes we examined experience SCE at a much higher rate than the rings examined by Gatti, *et al*. (1979). One of those was *R(1)2*, which, in our hands, also exhibited a low rate of bridge formation, making this explanation feasible.

The origin of SCU events cannot be discerned from our experiments. In previous work, Hinton moved the heterochromatin of an unstable ring chromosome to a linear chromosome, and found that the “linearized” ring still produced chromosome bridges and lethality. Since a SCE is not expected to cause a problem with a linear chromosome, Hinton concluded that SCU was the basis of anaphase bridges with the ring that he studied. How the break and rejoining might have arisen was not determined, other than it mapped to heterochromatin (Hinton 1957). In our experiments, the break that preceded SCU may have occurred in the previous anaphase, arising through breakage of an anaphase bridge. We previously showed that double bridges produced by SCE can break in anaphase, making this course of events feasible (Hill and Golic 2015). It is also conceivable that one of a pair of catenated sister chromatids might break as a means of resolving the bridge. These events might be followed by SCU in a subsequent cell cycle.

One additional conceptual problem with ring chromosomes (and linear chromosomes) is incomplete replication (Fig. 1F). The added topological problems presented by replicating a ring chromosome might make replication more difficult, resulting in delayed completion and anaphase bridges. Bridges such as shown in Fig. 4E could result from incomplete replication, with only a tiny bit of unreplicated DNA connecting daughter chromatids. However, this could also be an example of catenanes caught in the process of topoisomerase-mediated separation. To positively identify such events will require more than simple DAPI staining.

In corn and in humans carrying a ring chromosome, varied numbers and forms of the ring chromosome are often observed in mosaic condition within an individual (McClintock 1932; McClintock 1938; Levan 1956; Guilherme et al. 2011; Pristyazhnyuk and Menzorov 2018). A variant frequently seen in humans is a double-sized dicentric ring arising from SCE. In metaphase preparations these chromosomes consist of two double-sized sister chromatids, and therefore must have arisen through some combination of SCE and nondisjuction. We saw no such double-rings in our examinations of *R(1;Y), R(2)* or *R(3).* Furthermore, in the work decribed here, the vast majority of cells were euploid. In *R(1;Y)/R(1;Y)* larvae, cells with an extra copy of the ring, though few in number, were seen in most brains and a few brains had single cells with only one ring. In spite of much more extensive examination of *R(3)* metaphases, only a single example of a cell with chromosome *3* aneuploidy was seen. The paucity of variants seen in Drosophila could have many causes, such as a relatively infrequent SCE in flies relative to humans and corn (Crossen et al. 1977; Gatti et al. 1979; Raposo et al. 2004), or a preference for bridges to break rather than experience nondisjunction.

Even if cells with whole-chromosome aneuploidy are produced, selection likely plays a large part in their elimination. A single Drosophila chromosome represents a much larger fraction of the genome than a single chromosome in corn or in humans and whole chromosome aneuploidy is likely to be frequently cell-lethal. However, variation in small rings has been observed in Drosophila. Hilliker (Hilliker 1978) examined a much smaller ring, consisting of chromosome *2* heterochromatin. He saw cells possessing anywhere between 0-3 copies, consistent with the idea that ring chromosomes in *Drosophila*, as in corn or humans, can exhibit cell-to-cell variation in copy number, but that these cells only survive if they are not grossly aneuploid. The generation of gynandromorphs, by loss of a ring-*X* from cells with two *X* chromosomes, is a special case. When loss occurs prior to blastoderm those cells can undergo dosage compensation and survive normally.

In considering the frequent anaphase bridges seen with the *R(1;Y)* and *R(3)* chromosomes, a critical question is how can an organism grow and survive if over 50% of its cells are experiencing anaphase bridges? There are likely several possible reasons. First, the bridge frequencies shown in Table 2 represent a snapshot of cells in anaphase. If the occurrence of a bridge delays a cell’s departure from anaphase into G1, then cells with problems will be overrepresented in these squash preparations. Second, as the time-lapse videos of embryonic mitoses showed, the majority of bridges produced by sister chromatid catenanes are resolved, and daughter cells can proceed to the next cell cycle. Since sister chromatid catenanes form a high proportion of bridges, solving this problem alone likely accounts for a significant proportion of surviving cells.

Although mitotic problems were frequent with *R(1;Y)* and *R(3)*, they were relatively rare with *R(1)2.* If simply existing as a ring were the entire source of mitotic problems, *R(1)2* should also show a large fraction of mitoses with bridges. This difference might be attributable to size: *R(1;Y)11AX2* is roughly the same size as chromosome *R(3)* owing to the presence of the entire *X* euchromatin and a portion of its heterochromatin plus most of the *Y* chromosome, with *R(1;Y)6AX2* being slightly smaller. On the other hand, *R(1)2* is much smaller, containing only *X* euchromatin and heterochromatin. Chromosome size has been proposed to contribute to the extent of problems experienced by ring chromosomes (McClintock 1938): a large chromosome provides more opportunity for catenation of sisters and could experience a higher rate of SCE. However this may not be generally true (Kistenmacher and Punnett 1970). In *Drosophila,* a previous examination of single and double-sized ring-X chromosomes showed that the single ring experienced more SCE events than the double ring (Gatti et al. 1979). Instead, chromosome content might influence sister chromatid catenation and SCE. In normal linear chromosomes, persistence of sister catenations into mitosis appears to occur primarily in heterochromatin (Porter and Farr 2004; Mills et al. 2018; Chu et al. 2022). Moreover, SCE also occurs primarily in heterochromatin (Gatti *et al*. 1979), and the *R(1;Y)* and *R(3)* chromosomes both have significantly more heterochromatin than *R(1)2*. Alternatively, since *R(1)2* has been maintained in stock for nearly a century, there has been ample time to accumulate genetic variants that reduce its misbehavior.

Our most surprising finding was the discovery of homolog capture, where the ring maintains a connection with its linear homolog into mitosis, producing a chromosome bridge during anaphase. This connection is almost certainly a remnant of the mitotic pairing of homologs. We hypothesize that the homolog capture anaphase bridges represent topologically intertwined ring and linear homologs, with the linear wrapped several times so that neither topoisomerase nor tension is able to immediately resolve the problem in anaphase. This explanation would require that homologs wind around each other during the process of pairing. Direct observation of homologs in mitotic prophase certainly suggests that this is true. Although homologs with free ends might wind around each other starting at one or both ends, current evidence suggests that homolog pairing initiates at multiple sites along chromosomes (Fung et al. 1998; Viets et al. 2019). To deal with possible entanglements and to complete pairing, assistance from topoisomerase II may be required. In the complex nuclear arrangement of decondensed homologs and heterologs, chromosomes are likely to frequently pass across or through each other (Amoiridis et al. 2024). Passage of chromosomes through each other to facilitate pairing could easily generate twisted or braided homologs.

We directly tested whether topoisomerase II-like activity is involved in pairing by asking whether, in a ring/ring homozygote, the homologs become catenated. Our results showed that catenation of ring homologs is a frequent occurrence. This could occur through repeated rounds of homologous recombination in the same way that repeated rounds of SCE can catenate sister chromatids. But, given the rarity of mitotic recombination (reviewed by Ashburner et al. 2005 and results presented here) relative to catenation of homologs, this is extremely unlikely. Instead, we conclude that these catenanes are a result of topoisomerase II participation in the process of homolog pairing.

This raises the question: if topoisomerase II can catenate homologs during pairing, why is it unable to fully decatenate ring chromosomes as a cell approaches mitosis? Sister chromatids, even in normal linear chromosomes, are connected by catenation of DNA loops that emanate from the chromosome axes (Broderick and Niedzwiedz 2015), and which must be resolved during mitosis. In eukaryotes there is sufficient topoisomerase II to achieve this with great regularity. However, ring chromosomes are likely to experience additional catenations that result from the replication of a circular molecule and consequently require additional topoisomerase II activity. An organism that has evolved with linear chromosomes may have insufficient topoisomerase II to deal with this. In support, small ring chromosomes in *Schizosaccharomyces pombe* are unstable, but can be stabilized by additional topoisomerase II (Murakami et al. 1995). It is also notable that ring chromosome anaphase bridges in *Drosophila* have topoisomerase II localized to the center of the bridge, suggesting an attempt to disentangle sister chromatids (Ferree et al. 2014). Additionally, because homolog pairing occurs before replication (Csink and Henikoff 1998; Joyce et al. 2012), the amount of topoisomerase II needed for pairing, even for ring chromosomes, is probably substantially less than is required to separate sister rings and homologs in mitosis.

Finally, our observation of catenation of homologous chromosomes has significant implications for the mechanism of mitotic pairing in Drosophila. Williams *et al*. (Williams et al. 2014) knocked down topoisomerase II function in Drosophila cells and found that mitotic pairing was reduced. This established a pro-pairing role for topoisomerase II, but the nature of its contribution was unknown. Our observation that ring homologs become catenated during interphase provides an explanation: topoisomerase II passes double-strand DNA molecules across each other to achieve full pairing.

## Acknowledgements

This work was supported by grant R35 GM136389 to KGG from the National Institutes of Health. HJH was partially supported by training grant T32GM141848 from the National Institutes of Health. Drosophila stocks were obtained from the Bloomington Drosophila Stock Center at the University of Indiana, USA, and from the Kyoto Drosophila Stock Center at the Kyoto Institute of Technology, Japan.

